# A Guide to the Quantitative Proteomic Profiles of the Cancer Cell Line Encyclopedia

**DOI:** 10.1101/2020.02.03.932384

**Authors:** David P. Nusinow, Steven P. Gygi

**Affiliations:** Department of Cell Biology, Harvard Medical School, Boston, MA 02115, USA

## Abstract

We recently reported the quantitative proteomics of 375 samples as part of the Cancer Cell Line Encyclopedia (Nusinow et al., 2020). Mass spectrometry-based proteomics data is broadly unfamiliar to most biologists in our experience, resulting in questions from analysts about how to use the data. From the proteomics community there was interest about how we normalized the data, as the scope of this project was so much larger than what has been commonly available. This paper serves as a guide to the data set to answer these questions and acts as a supplement to the main manuscript. The first part addresses users of the data, describing the experimental design, interpretation of the values, and dealing with standard issues in proteomics like multiple protein isoforms per gene and missing values. The second part of the manuscript details how we arrived at the normalization procedure reported in the paper, including the diagnostics used to assess multiple normalization schemes.

## Introduction

Mass spectrometry-based proteomics has seen significant advances in sensitivity and quantitative accuracy throughout the last decade. One aspect of these improvements was the development of increasingly capable multiplexing reagents alleviating both missing measurements between samples and instrument time requirements. This enabled the regular production of experimental designs with many samples and the ability to produce datasets with tens to hundreds of them, several of which have now been published (Frejno et al., 2017; Gholami et al., 2013; Lapek et al., 2017; Mertins et al., 2016; Pozniak et al., 2016; Vasaikar et al., 2019; Zhang et al., 2014, 2016).

One widely used community resource is the Cancer Cell Line Encyclopedia (CCLE), which encompasses multiple omics-level experiments performed on a collection of nearly 1,000 cell lines (Barretina et al., 2012; Ghandi et al., 2019; Li et al., 2019). As the technology continued to mature it appeared possible to attempt quantitative proteome profiling as part of this project. Deep quantitative proteomics covering the majority of the expressed proteome has not achieved this sample size in another single experiment at the time of this writing, and at the outset sample sizes were more regularly on the order of less than 10. Thus, how to normalize proteomics data across hundreds of samples, even using the then state of the art TMT 10-plex reagent, was unknown.

This paper is meant to serve two purposes. The first is to describe the basics of the data and act as a primer for how to use the dataset. We hope that interested biologists, computational and otherwise, will find this useful, especially those with minimal experience with proteomics data. We provide advice on our experiences in using the data including ways to manage missing values during analysis.

The second purpose of this paper is to report the details of how the normalization for the CCLE proteomics data was settled on. This is meant to be a sort of appendix to the main paper, and act as a guide to the process of normalizing these data so that others can use the method or a similar one in their own work. All of the methods were documented in the main manuscript, but the pitfalls and wrong turns along the way were not. We hope that it will be useful to the proteomics community trying to produce similar large datasets to understand our own experience on this project.

## Experimental Design

As described in the manuscript (Nusinow et al., 2020), the experiments were performed in a multiplex setup using a reagent named Tandem Mass Tags (TMT, **Figure 1**) that, at the start of the project, allowed 10 samples to be run in parallel on the mass spectrometer (McAlister et al., 2012, 2014; Werner et al., 2012). Each sample in the data (the column names on supplemental table S2 of Nusinow et al., 2020) denotes which of the forty-two 10-plexes it was part of. In each 10-plex there were nine biological samples and one “bridge” sample that allowed normalization across plexes. This was a mixture of ten cell lines from the CCLE selected for maximal gene expression diversity that was prepared in a large batch at the beginning of the project, aliquotted out, and frozen until needed per batch. This bridge sample is not included in the normalized data, leaving only nine samples per 10-plex.

**Figure 1:**
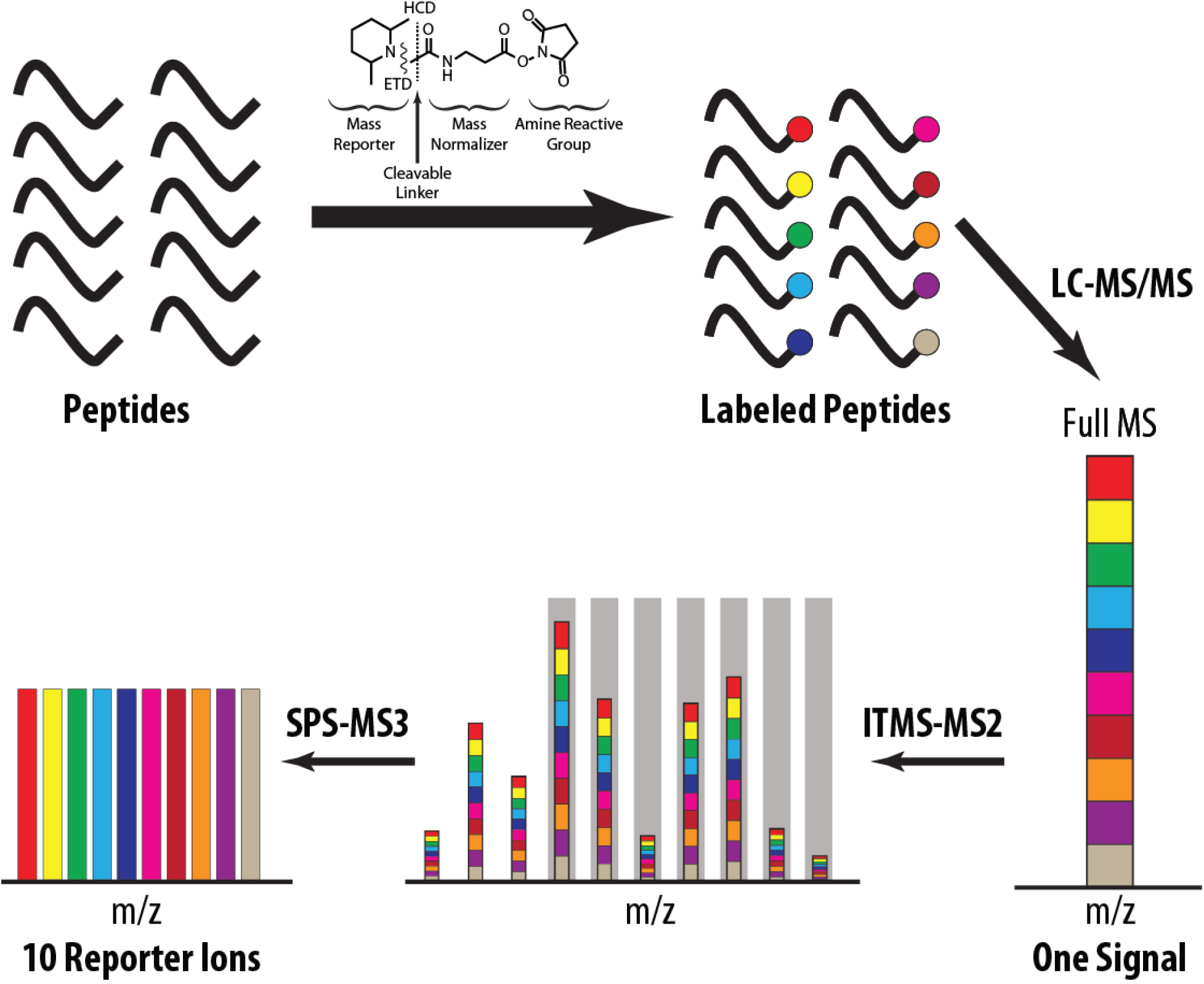
Multiplexed Proteome Analysis by TMT. Tryptic peptides were isolated from nine cell lines and the bridge sample and separately labeled with the TMT-10 reagents (top). Labeled peptides were combined and analyzed by LC-MS. The TMT reporter variants are isobaric, and thus coelute during chromatography and only show a single convoluted peak during MS analysis (bottom right) and multiple convoluted peaks during peptide sequencing via MS/MS (bottom middle). Upon MS2 fragment peak isolation for MS3 analysis (grey boxes in bottom middle) the TMT reporter fragments are deconvoluted and quantitative values can be read (bottom left).

In addition, three individual cell lines were repeated within the dataset. These lines have much poorer correlation between replicates than our biological replicate samples. We do not understand the cause of this but have left the samples in the data. If you choose to focus on only one replicate for those lines we recommend using the replicate with the best correlation to the RNASeq data, as this is our best quality control metric. From this diagnostic, SW948_LARGE_INTESTINE_TenPx20, CAL120_BREAST_TenPx28, and HCT15_LARGE_INTESTINE_TenPx18 should be the preferred versions for each cell line if you wish to choose only one.

A potential source of bias in the data is our inherent inability to fully block the experiment so as to randomize effects across plexes. We controlled for these things as well as possible and did not detect any biases in the data after normalization, but it is possible they are still there. Another possible source of bias is that samples with low protein yields were blocked to the latter plexes without disrupting tissue distribution among plexes. This led to decreased peptide yields but not proteins quantified. This can lead to less reliable quantitation among these samples, although in our experience with the SPS-MS3 method used here individual spectra routinely show high accuracy of quantitation.

## Interpreting the Normalized Data

A commonly asked question is what the quantitative values actually are. The normalization is detailed in Nusinow et al, 2020 and below, but the values are most closely log2-transformed ratios to the bridge sample in each plex. After taking this ratio, for each protein in each multiplex the values are mean centered so as to remove errors in measuring the bridge, but in most cases this does not shift the values much and they remain close to the log2 ratio to the bridge.

Importantly, these values are all relative values to the other values for that same protein and not absolute values. This means that comparing the levels of different proteins to each other without using something like a correlation to standardize values won’t produce meaningful results. Similarly, these values are not standardized, as in a z-transformation. Many data analysis methods suggest standardization before use but these data are not delivered that way, although it easy to standardize the values yourself.

A critical decision about this experiment is that the data are normalized to the total protein amount rather than cell numbers or per-protein mass. Thus, approximately equal amounts of total protein from lysate were processed between samples, but these will represent different numbers of cells depending on the cell line. Interpretation of any downstream analysis must keep this in mind.

## Using the Normalized Data

Details about the cell lines was provided as Supplementary Table S1 in Nusinow et al., 2020 and can also be found on the CCLE website. We used the internal CCLE nomenclature for cell line identification. DepMap has subsequently revised this ID scheme, and that project website provides annotation for cell lines that includes those IDs. Mapping between these sample IDs may be required for analyses of DepMap and future data sets.

The raw mass spectrometry data was searched using a database from UniProt (The UniProt Consortium, 2018), and all the annotations in the normalized data come from there including gene symbols. We include a Protein ID column, which is a unique identifier from UniProt that takes in to account isoform variations. Our protein counts are defined by these Protein IDs, so one gene symbol will often be represented by multiple Protein IDs because peptides belonging to different variants were quantified. Our data processing pipeline (Huttlin et al., 2010; Nusinow et al., 2020) reduces the number of protein IDs down to the minimal number that explain all of the identified isoforms, so there should not be more proteins identified than are necessary. Common ways to analyze the data are per-protein or collapsed down to per-gene symbol. In many cases, one isoform will be quantified in only some of the ten-plexes while another is quantified every time, so it could be beneficial to use the version quantified every time when collapsing down to gene symbols.

This raises another common issue with these data which is the preponderance of missing values. These are a standard problem in bottom-up proteomics that, in our experience, people outside the field are frequently unprepared to handle. Many out of the box software packages and algorithms don’t handle missing values well, so consideration is often needed in how to use those packages. Our approaches have been the following:

1. In cases where only complete values are allowed, such as in Principle Components Analysis, we use only those proteins that were quantified in every sample.
2. In cases where we compare proteins to each other, as in building a correlation network, we take only the values where measurements were taken in both cell lines and require a certain number of values to be present, usually arbitrarily set at 100 for global analyses.
3. In cases where we compare different samples within the same protein, we only take the non-missing values and require a certain number of values to be present in each subgroup and in total. This number depends on the sample size for the subgroups and is usually determined by running models with multiple test correction and inspecting the results manually.

A final note about missing values is that a common approach to handling them is imputation. While this is a popular approach, we feel that for a data set as complex as the CCLE the models underlying imputation will be far more error-prone than on experiments with a smaller number of instrumented variables. We have not evaluated imputation schemes on these data and urge caution if this is something you are interested in doing. We found that by using the approaches above many analyses are tractable while leaving intact the uncertainty in the missing values, which we find useful for having confidence in our results.

## Details of the Normalization

This portion of the manuscript details the work on normalizing the data. It is not meant for analysts who want to use the data to gain biological insights, but rather for computational proteomics researchers who are interested in the technical aspects of normalizing these kinds of data. It reproduces work presented in a poster at ASMS in 2018, which is also available upon request.

### The Approach

We began the initial work on normalizing the data when 6 ten-plexes, representing 54 samples, were completed, creating a proteomic dataset about as large as any then published. Following common methods for small sample numbers we initially used visual inspection of tissue clustering coherency in hierarchical clustering or PCA as the diagnostic for normalization effectiveness. We found very early after trying different normalization approaches that visual inspection of tissue clustering was intractable and that quantitative diagnostics for normalization were required. The diagnostics we developed for this study are described below.

Additionally, a parallel experiment in the Gygi lab to profile dozens of mouse liver proteomes (Chick et al., 2016) had found various technical artifacts by manual inspection that were also present in the CCLE data. Our ability to visually discover artifacts as well as the complexity of interpreting certain popular normalization schemes described below led us to prefer simple, readily interpretable normalization adjustments in preference to more complex ones. These adjustments always had to be targeted towards well understood defects in the data, and the adjustments were chosen based on our best understanding of the nature of those defects. While more complex methods were approximately as effective at removing defects based on our diagnostics, we chose to be conservative in order to better understand and control these already extremely complicated data.

### Normalization Diagnostics

As described above, the complexity of the data made it clear that quantitative diagnostics would be required to evaluate different normalization schemes. We knew a handful of things about the experiment that we leveraged to develop the diagnostics. Once we had completed the biological replicate samples, the correlations between them became valuable tools (**Figure 2A**).

**Figure 2:**
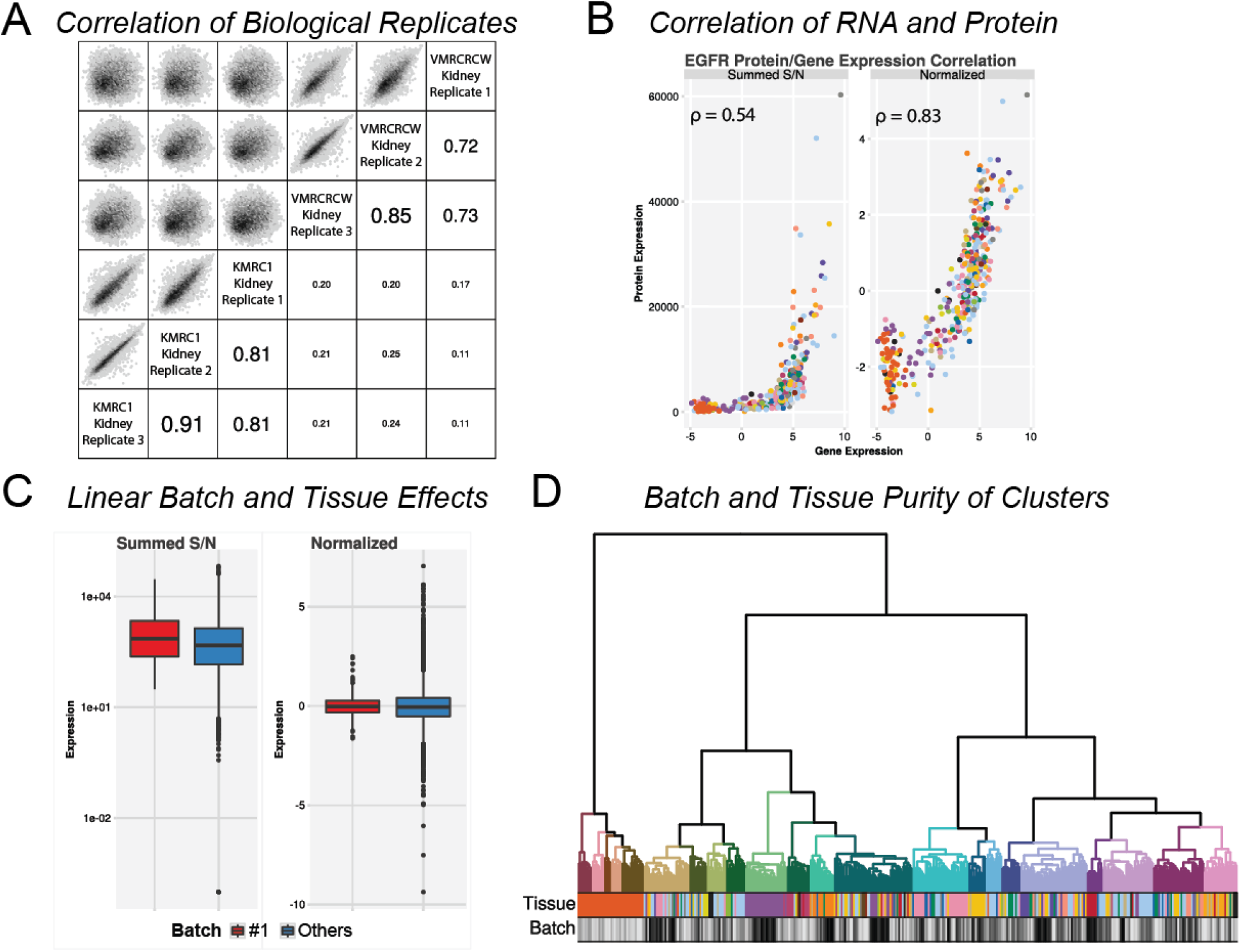
Diagnostics used to assess normalization schemes in this study. **(A)** Scatter plots of sample correlations between biological triplicates of two kidney-derived cell lines. All of the cell lines used in the first two ten-plexes were regrown and reprocessed after the initial set to act as a biological replicate control. Replicates correlate more than with the other cell line. Replicates 2 and 3 were grown together and processed a full year after replicate 1. **(B)** An example of correlated EGFR protein and RNA-Seq expression. Normalization helps the correlation, shown on each plot. **(C)** A motivating example for the linear modeling diagnostic. The distribution of expression of ten-plex batch #1 is shown compared to all other batches before and after normalization. The difference in mean values for batch and tissue effects can be quantified by linear modeling. **(D)** Hierarchical clustering of the full data set can be partitioned in to the same number of groups as there are tissues of origin in the sample set. Colored bars in the dendrogram denote clusters. Tissue and batch purity of the clusters can be quantified by Gini impurity.

Given the experimental design and the already existing data, other diagnostics were possible. The correlation between RNA and protein (**Figure 2B**) proved to be extremely useful in assessing both normalization and sample identity (Nusinow et al., 2020, Figure 2). Although RNA and protein do not correlate very well overall, we find that their correlation significantly improved with certain normalization procedures.

Another important diagnostic for us was linear modeling. We fit models on the entire protein data using dummy variables for the batch and tissue with no interactors. This approach was designed to uncover systematic shifts due to batch effects (**Figure 2C**). An important point here was that tissue effects were expected due to the underlying differences between tissues. Thus, the linear models were used to assess removal of batch effects by the batch coefficients and for accidental removal of real biology using the tissue coefficients. These types of linear model diagnostics are extremely adaptable to most experimental designs and do not require additional data like biological replicates or RNA expression data.

The final diagnostic we used was based on hierarchical clustering coherency of tissues and batches. Our expectation was that any normalization approach would attempt to minimize batch effects while maximizing tissue clustering, without expecting perfection for either. To quantify this effect, we used gini purity, related to the gini impurity scoring used in machine learning approaches such as classification trees (Breiman et al., 1984). One problem with this approach is that hierarchical clustering can be performed in any number of ways using different distance metrics and clustering approaches. We settled on one approach based on visual appearance and held that approach consistent while attempting different normalization methods. It might be possible to optimize the clustering approach to improve these diagnostics, but we are not aware of any good way to do this, and consider keeping the clustering method consistent to be the most important thing. Another problem with this approach is that it had low resolution. We found that the different normalization methods didn’t do much to change the clustering coherency, and even with the size of our data, the limited numbers of samples for some tissues meant that this diagnostic was often harder to interpret. We thus used it less than the other three approaches, but it provided a nicely visual method that encouraged us to keep it.

### Common Methods of Normalization

We initially looked at common normalization approaches from the gene expression field. Quantile normalization (Ritchie et al., 2015) is an approach that adjusts the position and shapes of the expression distributions per-sample to be consistent **(Figure 3A)**. After performing this method on the sample loading-adjusted protein data, a simple visual inspection by hierarchical clustering showed no tissue coherency and perfect batch coherency **(Figure 3B)**. We conclude that on these data quantile normalization exacerbates batch effects and should not be used.

**Figure 3:**
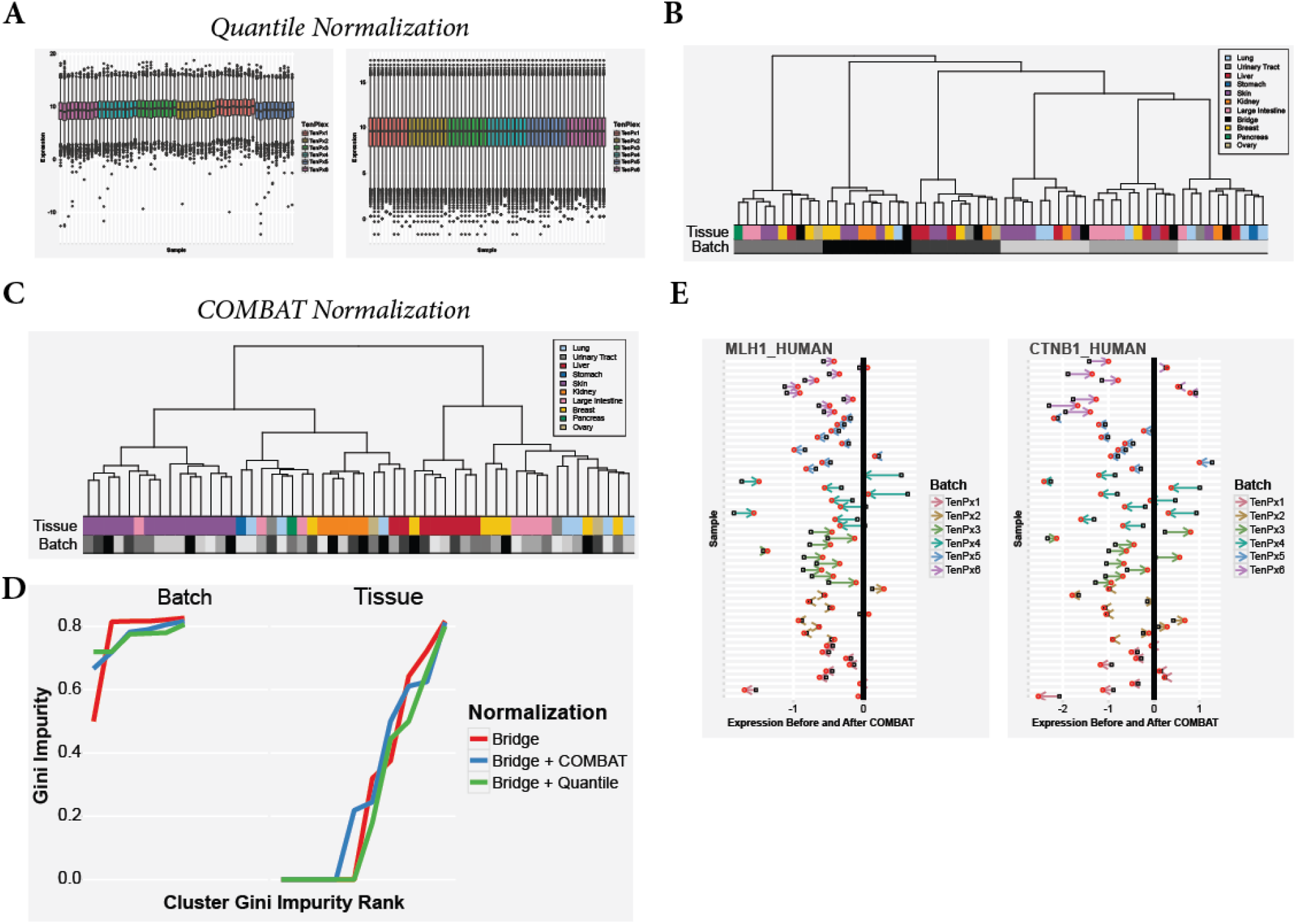
**(A-B)** The transformation of non-normalized expression distributions for the first six ten-plex groups **(A, left)** by quantile normalization **(A, right)**. **(B)** Hierarchical clustering of the first 6 ten-plex groups after quantile normalization creates an obvious batch effect (greyscale bars). **(C-E)** Hierarchical clustering after COMBAT normalization **(C)**, quantified by cluster Gini impurity **(D)**, shows no obvious batch effects and reasonably coherent tissue clustering. **(E)** Two examples of the effects of COMBAT normalization. COMBAT induces a complicated transformation resulting in overall compression of sample variance. The compression of individual samples is inconsistent within batches, with larger adjustments sometimes happening to more extreme values and sometimes to less extreme.

An extremely interesting approach is the COMBAT algorithm (Johnson et al., 2007). This method is an Empirical Bayes approach, which estimates prior distribution parameters from the entire data and then uses that prior to moderate large deviations from global averages. The standard implementation of COMBAT does not handle data with missing values, but we modified the implementation to allow for them as the underlying approach appears to be compatible with missing values. Testing COMBAT on the first 54 samples showed no batch effect and good tissue coherency by hierarchical clustering **(Figure 3C-D)** demonstrating that by global measures COMBAT performs well on proteomics data and appears compatible with missing values. Investigation of the effects of COMBAT on individual protein levels was complicated **(Figure 3E)**. Per sample effects were unpredictable, where in some cases extreme values shifted towards the mean a great deal while in other cases they hardly moved at all. Because of the challenges in interpreting the output of this normalization and the comparable performance of the approaches we developed later we did not pursue COMBAT further for this study, however it remains a compelling approach that merits further work.

### Normalizing for Total Protein Amount

A standard approach in proteomics is to load consistent total protein amounts per sample, which is the approach we used here. After labeling we performed “ratio checks” on a small portion of unfractionated sample before recombining the full amount, and adjusting the protein loading if the deviations for any sample were too high. Even with these adjustments, subsequent normalization of the full data for sample loading differences has to be performed. This can be done in several ways, but the most common that we are aware of are to adjust based on the mean, median, and total summed value of all the reporter values for all proteins in a sample. We choose to perform this normalization first as a standard part of TMT-based protein expression profiling in our pipeline because it is so commonly needed.

We checked the three different adjustment methods using our diagnostics **(Figure 4)**. The column mean and sum showed no differences at all **(Figure 4)**. Using our linear modeling diagnostics, normalizing based on the sample mean or sum showed a residual batch effect remaining after normalization, but also a preservation of tissue effects **(Figure 4A)**. Normalizing based on the sample median should protect against highly deviant batches and samples better than the mean, and indeed this showed a lesser residual batch effect than sum or mean normalization **(Figure 4A)**. However, along with the reduction in batch effect was a reduction in tissue effect with the median **(Figure 4A)**. These effects were also seen based on clustering coherence **(Figure 4B)**. There was a slightly better correlation between protein and gene expression using mean versus median normalization, with 0.422 vs 0.420 average Pearson correlation for mean and median adjustments respectively. The difference was indistinguishable globally though (data not shown).

**Figure 4:**
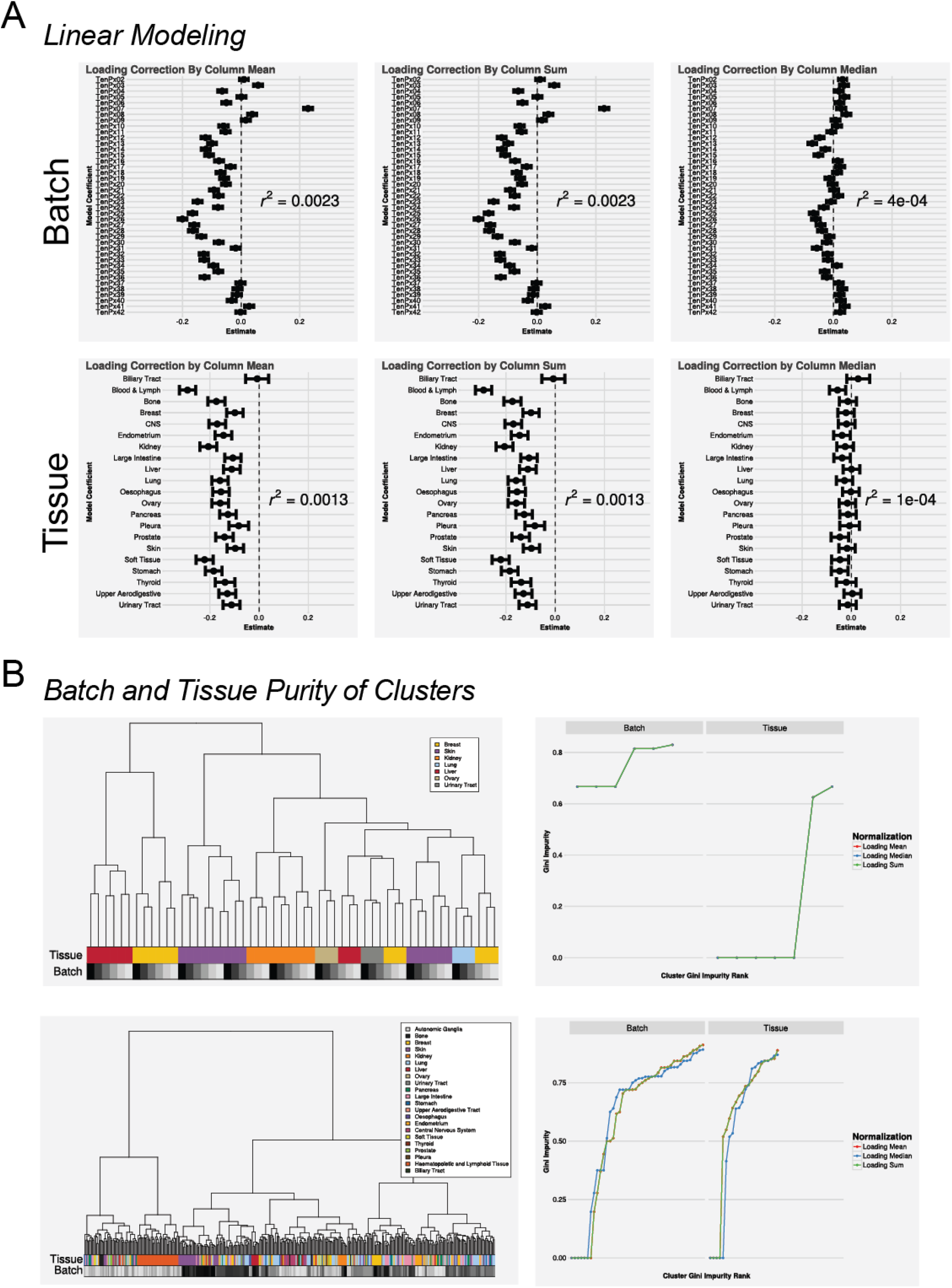
Adjustment for Differential Protein Loading. Sample loading adjustments using the per-sample column sum, mean, and median were tested across multiple diagnostics. **(A)** Linear model coefficient plots for batch (top row) and tissue (bottom row), along with associated r^2^ values for the model. Columnwise sum and mean shifts result in indistinguishable models. The median shift corrects more batch effects at the cost of removing tissue-based information. **(B)** Hierarchical clustering (left) and associated cluster Gini impurity values (right) for biological replicates (top) and the full data set (bottom). All loading adjustments result in equivalent clustering of the replicate samples. As in the linear models in (A) the median correction removes slightly more batch effects but performs worse on tissue-based effects than the sum and mean adjustments.

Although the median-based sample loading correction was better at removing the batch effects than the sum or mean, the removal of the tissue effect indicated that it was also removing important biological signal. Indeed, samples with large numbers of outlier points are likely due to different cellular identities based on factors like the tissue of origin. Because of the complexity of the CCLE, it appears that a sum or mean-based loading normalization is appropriate, however in experiments with fewer distinct cellular identities the median-based approach might be more appropriate.

### Normalizing for Per-Protein Batch Effects

The residual batch effect following sample normalization by the column sum required investigation. Visual inspection of the data found proteins where all of the values in a single multiplexed experiment was shifted. An extreme example is shown in **Figure 5A**, but many less extreme examples were clear. Because the samples were blocked attempting to evenly distribute the tissues of origin with random individual selection as well as was possible, our expectation was that there should be no cases where a protein happens to be consistently up- or downregulated in all 9 samples in a ten-plex. While this is formally possible, it is unlikely.

**Figure 5:**
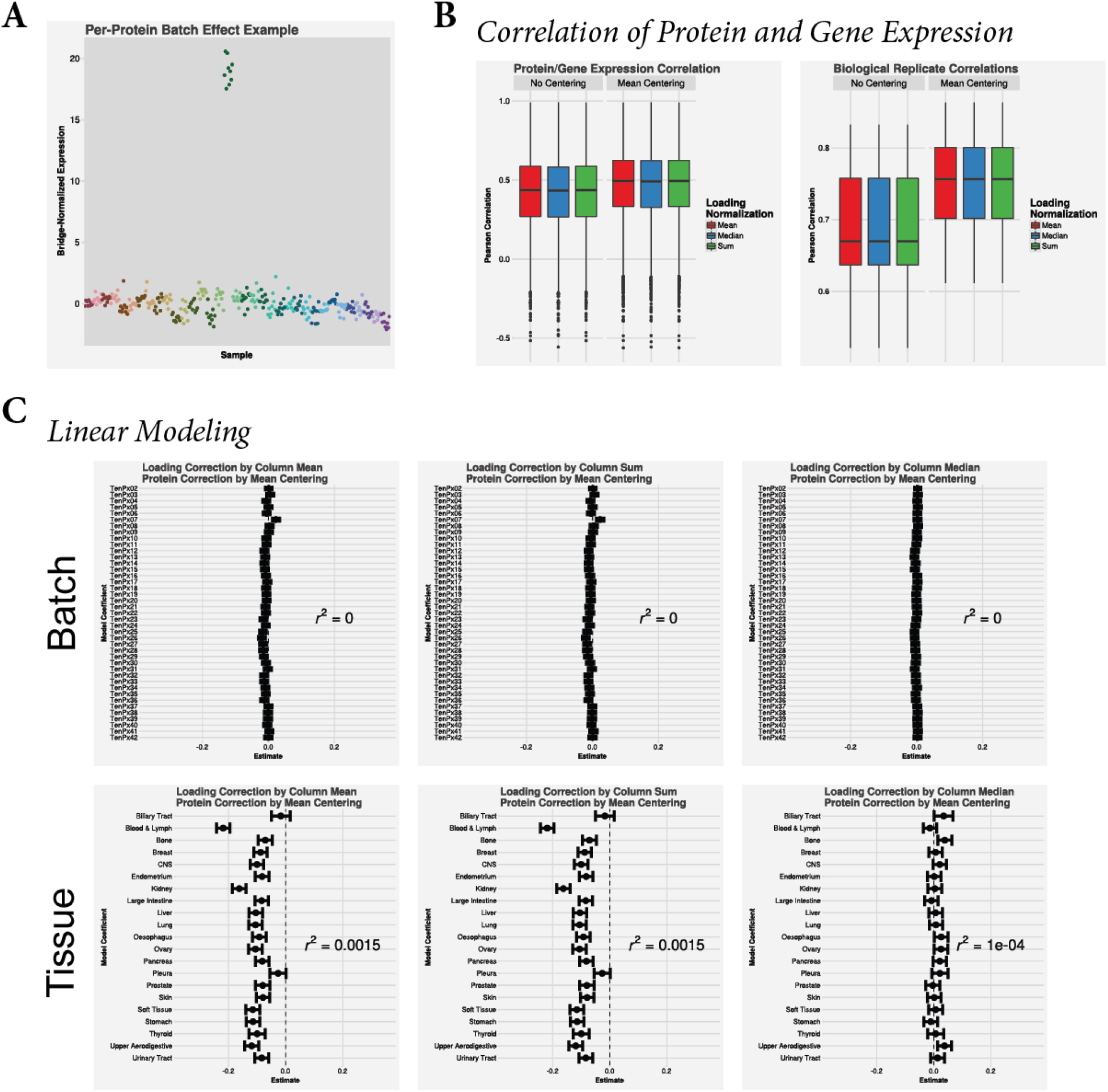
Per-Protein Batch Effect Normalization. **(A)** An example of the remaining artifacts we observed following loading and bridge normalization. This protein shows a large increase in the normalized levels in one ten-plex. This is likely an artifact. We correct this via mean centering each set of proteins within a batch after the preceding normalization steps. **(B)** This correction significantly improved both the correlation to gene expression in the full data set (left) and the correlation between biological replicates (right). **(C)** Linear model coefficient plots for both batch (top row) and tissue (bottom row). The batch effect still present after the previous correction has been reduced dramatically. The tissue-based effect remains after per-protein mean centering.

What is more likely is that there is some random error in the measurement of the bridge sample causing a systematic shift in the other measurements in that group. If this is the case, our model is that, on average, all the different 10-plex experiments will have the same average expression. Thus, our normalization to correct for per-protein bridge measurement errors is to enforce the same average expression by shifting all the expression values in that plex by the difference between that average expression and 0, making all values center on 0. This is referred to as mean-centering.

The effects of this on the biological replicate correlation and the RNA/protein correlations are shown in **Figure 5B**. In both cases, mean-centering the values improved the correlations over normalizing to the bridge alone. The linear modeling results showed that this normalization removed the residual batch effects from sum or mean normalizing for sample loading **(Figure 5C)** and also improved the batch effect removal from the median normalizing for sample loading. However, following sum or mean adjustments for sample loading, per-protein per-plex mean-centering also retained the tissue information. In contrast, it had been removed and was not recovered after median-based loading adjustment **(Figure 5C)**.

It is possible to median-center per-protein per-plex instead of mean-centering. We did attempt this and found that it performed less well than mean-centering on our global diagnostics (data not shown). A brief investigation indicated that, while this did protect against individual outlier samples, most measurements did not contain these outliers. In those cases, the estimate provided by the median based on only 9 samples was not as good as that of the mean. As multiplexed proteomics improves, perhaps even in the case of the current 16-plex reagent, a median-based adjustment might be better, but in this case the mean was superior.

An alternative experimental design uses two bridge channels and takes an average between them to standardize each protein within a plex (Lapek et al., 2017; Plubell et al., 2017). These methods were unpublished until we were well in to our study, but are designed to alleviate the issues with improper bridge measurement that we correct in the CCLE through mean centering. For an experiment with as complex a design as the CCLE it is unclear whether the increased number of plexes and the accompanying batch effects are worth the tradeoff of the residual error after mean centering. For simpler experimental designs, especially those without the expectation that the average protein expression would be centered (e.g. time series) these approaches might be more suited and should be considered when designing the experiment.

### Conclusions

In **Figure 6** we show the stepwise process of normalizing the expression of a single protein where the improvements to the data are visible at each step, until there are no obvious visible batch effects. Plots like these and the diagnostics we described gave us confidence in the normalization approach. We did investigate other minor differences like reordering the steps, but in the cases we tried there were no clear improvements.

**Figure 6:**
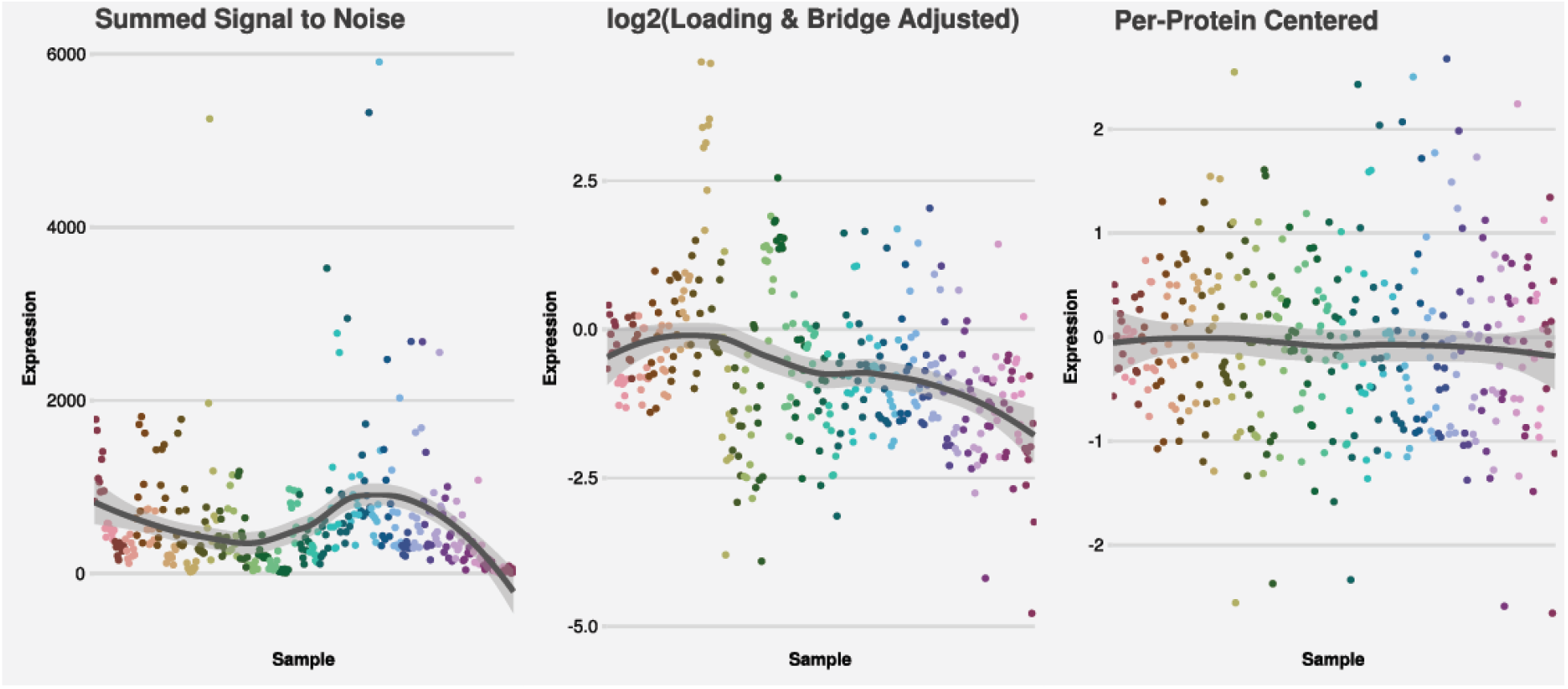
Demonstration of the stepwise effects of the normalization protocol used in this study. The stepwise effects of the normalization approach is shown above for the protein PIK3R1. Samples are colored by batch. Batch effects are progressively removed at each step, resulting in an expression profile lacking any obvious batch effect.

The key point we took away from this exercise is to have good quantitative diagnostics for normalizing the data based on the experimental design. We were fortunate to have resources to do biological replicates and had available gene expression data, which helped us immensely, but the basic linear modeling approaches are fully adaptable to other experimental designs without those resources.

No normalization procedure will fit all data because normalization is inherently tied to the defects and assumptions built in to a given experiment’s design. Indeed, we have seen this normalization procedure fail on other experiments in the lab with different designs and constraints. Good diagnostics and visualization tools are key to developing the right normalization strategy for any experiment, and prove invaluable for assessing schemes from the simple to the complex.

## References

Barretina, J., Caponigro, G., Stransky, N., Venkatesan, K., Margolin, A.A., Kim, S., Wilson, C.J., Lehár, J., Kryukov, G.V., Sonkin, D., et al. (2012). The Cancer Cell Line Encyclopedia enables predictive modelling of anticancer drug sensitivity. Nature 483, 603–307.

Breiman, L., Friedman, J., Stone, C.J., and Olshen, R.A. (1984). Classification and Regression Trees (Taylor & Francis).

Chick, J.M., Munger, S.C., Simecek, P., Huttlin, E.L., Choi, K., Gatti, D.M., Raghupathy, N., Svenson, K.L., Churchill, G.A., and Gygi, S.P. (2016). Defining the consequences of genetic variation on a proteome-wide scale. Nature 534, 500–505.

Frejno, M., Chiozzi, R.Z., Wilhelm, M., Koch, H., Zheng, R., Klaeger, S., Ruprecht, B., Meng, C., Kramer, K., Jarzab, A., et al. (2017). Pharmacoproteomic characterisation of human colon and rectal cancer. Mol. Syst. Biol. 13, 951.

Ghandi, M., Huang, F.W., Jané-Valbuena, J., Kryukov, G.V., Lo, C.C., McDonald, E.R., Barretina, J., Gelfand, E.T., Bielski, C.M., Li, H., et al. (2019). Next-generation characterization of the Cancer Cell Line Encyclopedia. Nature 569, 503.

Gholami, A.M., Hahne, H., Wu, Z., Auer, F.J., Meng, C., Wilhelm, M., and Kuster, B. (2013). Global Proteome Analysis of the NCI-60 Cell Line Panel. Cell Rep. 4, 609–620.

Huttlin, E.L., Jedrychowski, M.P., Elias, J.E., Goswami, T., Rad, R., Beausoleil, S.A., Villén, J., Haas, W., Sowa, M.E., and Gygi, S.P. (2010). A Tissue-Specific Atlas of Mouse Protein Phosphorylation and Expression. Cell 143, 1174–1189.

Johnson, W.E., Li, C., and Rabinovic, A. (2007). Adjusting batch effects in microarray expression data using empirical Bayes methods. Biostat. Oxf. Engl. 8, 118–127.

Lapek, J.D., Greninger, P., Morris, R., Amzallag, A., Pruteanu-Malinici, I., Benes, C.H., and Haas, W. (2017). Detection of dysregulated protein-association networks by high-throughput proteomics predicts cancer vulnerabilities. Nat. Biotechnol.

Li, H., Ning, S., Ghandi, M., Kryukov, G.V., Gopal, S., Deik, A., Souza, A., Pierce, K., Keskula, P., Hernandez, D., et al. (2019). The landscape of cancer cell line metabolism. Nat. Med. 25, 850–860.

McAlister, G.C., Huttlin, E.L., Haas, W., Ting, L., Jedrychowski, M.P., Rogers, J.C., Kuhn, K., Pike, I., Grothe, R.A., Blethrow, J.D., et al. (2012). Increasing the Multiplexing Capacity of TMTs Using Reporter Ion Isotopologues with Isobaric Masses. Anal. Chem. 84, 7469–7478.

McAlister, G.C., Nusinow, D.P., Jedrychowski, M.P., Wühr, M., Huttlin, E.L., Erickson, B.K., Rad, R., Haas, W., and Gygi, S.P. (2014). MultiNotch MS3 Enables Accurate, Sensitive, and Multiplexed Detection of Differential Expression across Cancer Cell Line Proteomes. Anal. Chem. 86, 7150–7158.

Mertins, P., Mani, D.R., Ruggles, K.V., Gillette, M.A., Clauser, K.R., Wang, P., Wang, X., Qiao, J.W., Cao, S., Petralia, F., et al. (2016). Proteogenomics connects somatic mutations to signalling in breast cancer. Nature 534, 55–62.

Nusinow, D.P., Szpyt, J., Ghandi, M., Rose, C.M., McDonald, E.R., Kalocsay, M., Jané-Valbuena, J., Gelfand, E., Schweppe, D.K., Jedrychowski, M., et al. (2020). Quantitative Proteomics of the Cancer Cell Line Encyclopedia. Cell 180, 387–402.e16.

Plubell, D.L., Wilmarth, P.A., Zhao, Y., Fenton, A.M., Minnier, J., Reddy, A.P., Klimek, J., Yang, X., David, L.L., and Pamir, N. (2017). Extended Multiplexing of Tandem Mass Tags (TMT) Labeling Reveals Age and High Fat Diet Specific Proteome Changes in Mouse Epididymal Adipose Tissue. Mol. Cell. Proteomics 16, 873–890.

Pozniak, Y., Balint-Lahat, N., Rudolph, J.D., Lindskog, C., Katzir, R., Avivi, C., Pontén, F., Ruppin, E., Barshack, I., and Geiger, T. (2016). System-wide Clinical Proteomics of Breast Cancer Reveals Global Remodeling of Tissue Homeostasis. Cell Syst. 2, 172–184.

Ritchie, M.E., Phipson, B., Wu, D., Hu, Y., Law, C.W., Shi, W., and Smyth, G.K. (2015). limma powers differential expression analyses for RNA-sequencing and microarray studies. Nucleic Acids Res. 43, e47.

The UniProt Consortium (2018). UniProt: the universal protein knowledgebase. Nucleic Acids Res. 46, 2699–2699.

Vasaikar, S., Huang, C., Wang, X., Petyuk, V.A., Savage, S.R., Wen, B., Dou, Y., Zhang, Y., Shi, Z., Arshad, O.A., et al. (2019). Proteogenomic Analysis of Human Colon Cancer Reveals New Therapeutic Opportunities. Cell 177, 1035–1049.e19.

Werner, T., Becher, I., Sweetman, G., Doce, C., Savitski, M.M., and Bantscheff, M. (2012). High-Resolution Enabled TMT 8-plexing. Anal. Chem. 84, 7188–7194.

Zhang, B., Wang, J., Wang, X., Zhu, J., Liu, Q., Shi, Z., Chambers, M.C., Zimmerman, L.J., Shaddox, K.F., Kim, S., et al. (2014). Proteogenomic characterization of human colon and rectal cancer. Nature advance online publication.

Zhang, H., Liu, T., Zhang, Z., Payne, S.H., Zhang, B., McDermott, J.E., Zhou, J.-Y., Petyuk, V.A., Chen, L., Ray, D., et al. (2016). Integrated Proteogenomic Characterization of Human High-Grade Serous Ovarian Cancer. Cell 0.

